# Mixture model investigation of the inner-outer asymmetry in visual crowding reveals a heavier weight towards the visual periphery

**DOI:** 10.1101/2020.06.07.138412

**Authors:** Adi Shechter, Amit Yashar

## Abstract

Crowding, the failure to identify a peripheral item in clutter, is an essential bottleneck in visual information processing. A hallmark characteristic of crowding is the inner-outer asymmetry in which the outer flanker (more eccentric) produces stronger interference than the inner one (closer to the fovea). We tested the contribution of the inner-outer asymmetry to the pattern of crowding errors in a typical radial crowding display in which both flankers are presented simultaneously on the horizontal meridian. In two experiments, observers were asked to estimate the orientation of a Gabor target. Instead of the target, observers reported the outer flanker much more frequently than the inner one. When the target was the outer Gabor, crowding was reduced. Furthermore, when there were four flankers, two on each side of the target, observers misreported the outer flanker adjacent to the target, not the outermost flanker. Model comparisons suggested that orientation crowding reflects sampling over a weighted sum of the represented features, in which the outer flanker is more heavily weighted compared to the inner one. Our findings reveal a counterintuitive phenomenon: in a radial arrangement of orientation crowding, within a region of selection, the outer item dominates appearance more than the inner one.

## Introduction

Crowding refers to our inability to identify an object (primarily in the peripheral visual field) because it is presented with nearby objects^1–6^. Crowding is considered to be an impediment to reading^7^, face recognition^8^, eye movements^9,10^, visual search^3^ as well as deficits like macular degeneration^11^, amblyopia^12^ and dyslexia^13^. Research has characterized crowding and distinguished it from other spatial interferences, such as masking^6,14^, lateral interaction^15^ and surround suppression^16^. Yet, the underlying processes of crowding are still unclear^3,5,17,18^. Researchers have associated crowding with a variety of factors: the reduction in spatial resolution in the periphery due to increased receptive field size of cells at early visual areas^19–21^, decreased cortical magnification^22^ and reduced attentional resolution^23–26^.

A hallmark characteristic of crowding is the inner-outer asymmetry: an outer flanker (more peripheral) produces stronger interference than an inner one (closer to the fovea)^1,4,7–9,27–33^ (but see^36,37^). The asymmetry effect has been demonstrated with different stimulus types and tasks, such as letter recognition^36^, face recognition^28^ and Gabor orientation discrimination^16^. Nevertheless, the mechanisms underlying this phenomenon remain a matter of debate.

Researchers have proposed various causes of the inner-outer asymmetry. According to the cortical magnification account, the smaller cortical distance between the outer flanker and the target leads to stronger interference between these two items^22,37^. Other accounts postulate that increased receptive field size in the periphery biases sampling rate toward the outer flanker^27^ or leads to sparse selection in the visual periphery ^31^. Finally, the attentional account attributes display layout and experimental task in biasing spatial attention outwards, leading to the selection of the outer flanker^23,24^. Yet, none of the proposed accounts directly link the inner-outer asymmetry with recent models of crowding.

Proposed models of crowding were often classified to either pooling models^6,38–43^ or substitution models^18,44–46^. According to pooling models, under crowding, due to inappropriate integration field size in the periphery, observers simultaneously detect and pool excessive information of low-level features, including those that belong to the flankers^6,20,38–41^. In the more simple form of pooling models, crowding reflects an averaging of target and flanker features. For example, simple averaging errors were observed in judging target orientation^41^ and target position^38^. However, simple pooling models cannot explain the well-documented finding showing that under crowding conditions observers often misreport a flanker as the target^18,44–46^. Such misreport errors led to the proposal of substitution models, according to which increased location uncertainty in the periphery^47–49^ renders observers failure to spatially differentiate between the target and the flankers.

In recent years, researchers were able to reconcile these conflicting findings by proposing more advanced pooling models that can predict both averaging and misreporting errors (reviewed by Rosenholtz et al^50^). For example, several studies have proposed a population coding model that codes target and flanker features as a weighted sum within a receptive field and can explain both averaging and misreporting errors of orientation^39,42,51^, color and spatial frequency^51^. Investigations of these models often use estimation reports in which observers estimated the target feature in a continuous space; for example, reporting the orientation of the target by adjusting the orientation of a probe. In such tasks, observers often reported the flanker instead of the target^44,51,52^. By fitting a probabilistic mixture model with a misreport component (probability of reporting a flanker value), researchers were able to independently assess the precision of reported item and the contribution of the flankers to the distribution of the estimation errors^44,51^. This *misreport* mixture model acts similarly to the population coding model and can, therefore, serve to determine the relative activation of each item in the population^51^. However, whether the relative contribution, and hence activation, of the outer flanker is different from the inner one is still unclear.

Here, we addressed this issue by investigating the inner-outer asymmetry in a radial Gabor orientation crowding display with an orientation estimation task. We separately assessed the contribution of the inner and outer flankers to the pattern of crowding errors by comparing between various mixture models. Our findings show that a misreport mixture with a separate misreport component for the inner and the outer flanker outperformed all other tested models. Interestingly, we revealed that the misreport rate was much higher for the outer flanker than the inner one.

## Experiment 1

Observers performed an orientation estimation task of peripheral sinusoidal gratings (Gabor patches) in which the target (7° eccentricity) appeared alone (uncrowded conditions) or flanked (crowded conditions) by either two (two-flanker condition) or four (four-flanker condition) Gabors. Target and flankers were arranged on the horizontal meridian, either to the left or to the right of fixation. The centre-to-centre distance between two adjacent items was 1.5°. In crowded conditions, the target was always in the middle of the string of Gabors, such that in the two-flanker condition there was one inner flanker and one outer flanker, whereas in the four-flanker condition there were two inner flankers and two outer flankers with respect to the target (**Fig. 1A & B**).

**Figure 1.**
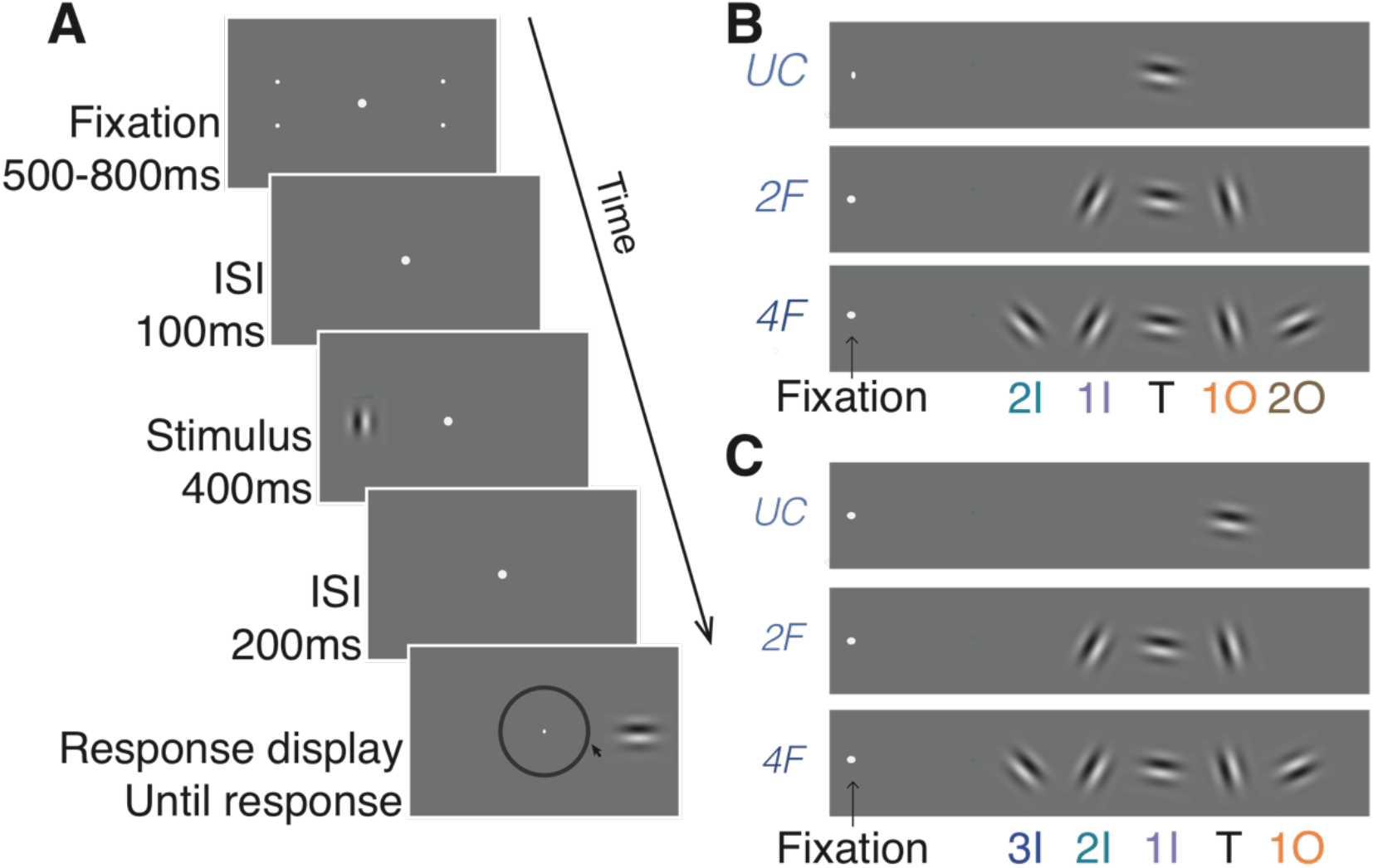
Stimulus type and crowding conditions. (**A**) Illustration of the sequence of events within a trial. During the fixation display, two pairs of dots, one pair in each hemifield, indicated the two possible locations (eccentricity) of the target. Observers estimated the orientation of the Gabor target by pointing the mouse cursor on a wheel. (**B**) The three stimulus conditions in Experiment 1. In Experiment 1 the target was always the intermediate Gabor, whereas in (**C**) Experiment 2, the target was the Gabor located one position out from the intermediate position. 3I: 3^rd^ inner flanker, 2I: 2^nd^ inner flanker, 1I: 1^st^ inner flanker, T: target, 1O: 1^st^ outer flanker, 2O: 2^nd^ outer flanker.

### Method

#### Observers

Thirteen undergraduate and graduate students (11 females: age range= 20 - 31 years, M=25.23, SD=3.32) from The University of Haifa participated in this experiment for course credit or a 40 ILS per hour monetary payment (around $12). On the basis of a priori power analysis, using effect sizes from previous studies^51^, we estimated that a sample size of 12 observers was required to detect a crowding effect with 95% power, given a .05 significance criterion. We collected data from one more observer in anticipation of possible dropouts or equipment failure. All observers were naive to the purposes of the experiment. All observers reported having normal or corrected-to-normal visual acuity and normal colour vision, and none reported attention deficits or epilepsy. The observers signed a written informed consent form before the experiment. This study was carried out in accordance with the relevant guidelines and regulations. The University Committee on Activities Involving Human Subjects at The University of Haifa approved the experimental procedures (No. 373/18).

#### Apparatus

Stimuli were programmed in Matlab (The MathWorks, Inc., Natick, MA) with the Psychophysics Toolbox extensions^53^ and presented on a gamma-corrected 21-in CRT monitor (SGI, with 1280 × 960 resolution and 85-Hz refresh rate) connected with an IMac. Eye movements were monitored with an Eyelink 1000 Plus (SR Research, Ottawa, ON, Canada), by using a chin rest at a 57-cm viewing distance. Observers reported their responses by using the mouse.

#### Stimuli and procedure

**Figure 1A** and **B** illustrate trial sequence and stimulus conditions in Experiment 1. All stimuli were presented on a grey background (56 cd/m^2^). Each trial began with the presentation of a fixation display. The fixation display consisted of a fixation mark (a centred dot subtending 0.1°) along with two peripheral pairs of place holders, one pair in the right hemifield and one in the left hemifield (7° eccentricity and ±2° vertical offset from the horizontal meridian), which indicated the eccentricity of the upcoming target. Both fixation mark and place holders were white (112 cd/m^2^).

Following observers’ fixation for a random duration lasting between 500 and 800 ms, the fixation mark appeared for an interstimulus interval (ISI) of 100ms. The stimulus display appeared for 400ms. The stimulus display consisted of the fixation mark along with the target; a sinusoidal grating (Gabors) with a 2D Gaussian spatial envelope (standard deviation 0.55° and 85% contrast); and a spatial frequency of 1.5 cycles per degree (cpd). The target appeared on the horizontal meridian with 7° eccentricity, either in the right or left hemifield. The target could appear alone (uncrowded condition) or along with two flankers (two-flanker crowded condition) or four flankers (four-flanker crowded condition), which were located on the horizontal meridian on either side of the target. The centre-to-centre distance between two adjacent items was 1.5°. In the two-flanker condition, flankers’ eccentricities were 5.5° and 8.5° for the first inner (1I) and first outer (1O) flankers respectively. In the four-flanker condition, flankers’ eccentricities were 4°, 5.5°, 8.5° and 10° for the second inner (2I), first inner (1I), first outer (1O) and second outer (2O) flankers respectively. Thus, all flankers appeared within the window of crowding (< 0.5 of eccentricity). Target and flanker orientation were selected at random from a circular parameter space of 180 values evenly distributed between 1° and 180°, with the restriction that each of the flanker’s orientation differed from that of the target by at least 15°.

Following the stimulus display, an additional fixation screen was presented for 200ms, which was followed by a response display. During the response display, observers were required to use a mouse cursor to adjust the probe Gabor’s orientation to match the orientation of the target by selecting a position on the orientation wheel (marked by a black circle 0.08° thick with an inner radius of 3.8° at the centre of the screen). Following an observer’s response, a blank inter trial interval (ITI) appeared for 400ms. In each trial, we monitored eye fixation using an eye tracker (see Apparatus). Trials in which fixation was broken (>1.5° from fixation mark) were terminated and rerun at the end of the block.

#### Design

Each observer completed 15 blocks of 40 trials (600 trials in total) over a 60-min session. There were 200 trials in each of the three crowding conditions (uncrowded condition, two-flanker condition and four-flanker condition). Crowding conditions were randomly mixed within each block. To reduce location uncertainty, observers were told that the pairs of dots have the same eccentricity as the upcoming target. The experiment began with two practice blocks of 10 trials. Observers were encouraged to take a short rest between blocks.

##### Models and analyses

For each trial, we calculated the estimation error for orientation by subtracting the true value of the target from the estimation value, such that zero indicates target value. To assess the contribution of the flankers to the error distribution (i.e. models with misreport components) we calculated flankers’ values by subtracting the true value of the target from the values of the flankers. We analyzed the error distributions by fitting probabilistic mixture models, which were developed from the standard model as well as the standard with misreport model^29^. We compared among four models:

*The standard mixture model* (Equation 1), as described in Yashar et al^51^, has two components: a von Mises (circular) distribution, which describes the probability density of reports around the target’s orientation, and a uniform distribution, which describes the probability of reports unrelated to the target (guessing rate). The model has two free parameters (γ, σ):

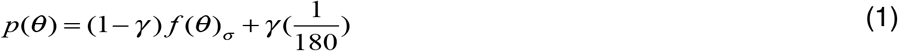

where θ is the value of the estimation error, γ is the proportion of trials in which participants are randomly guessing (guessing rate), and *f*(θ)_σ_ is the von Mises distribution with a standard deviation (variability) σ (the mean is set to zero), and *n* is the total number of possible values for the target’s feature.

*The standard misreport model* (Equation 2) adds a misreport component to the standard mixture model, which describes the probability of reporting one of the flankers to be the target. The model has three free parameters (γ, σ, β):

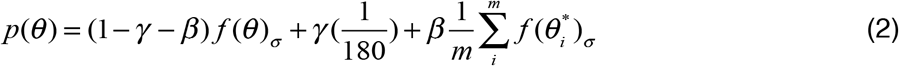

where β is the probability of reporting a flanker as the target, *m* is the total number of flankers (two or four in the present study), and *f*(θ*)_*i*_ is the von Mises distribution around the feature of the *i* flanker. Notice that the von Mises distribution of the estimation errors *f* (*θ*) describes the distribution when the observer correctly estimated the target’s feature, thus its mean is zero; whereas, for the distribution of misreporting one flanker as the target (*f*(*θ*\*)), the mean would be the feature distance of the corresponding flanker to that of the target. The variability of the distributions for each stimulus was assumed to be the same.

To test for sampling bias over the flankers (differences in sampling weights across flankers), we also fitted models with an independent β component for each flanker.

*The two-misreport model* (Equation 3) has a separate misreport component for each of the flankers in the two-flanker condition, one for the inner flanker and one for the outer flanker. This model has four free parameters (γ, θ, *β*_1*I*_, *β*_1*O*_):

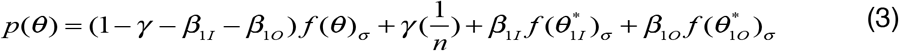

where β values represent the weights for each flankers: *β*_1*I*_ is the probability of misreporting the inner flanker as the target, and *β*_1*O*_ is the probability of misreporting the outer flanker as the target.

*The four-misreport mode*l (Equation 4) has a separate misreport component for each of the flankers in the four-flanker condition. The model has six free parameters (γ, θ, *β*_1*I*_, *β*_1*O*_, *β*_2*I*_, *β*_2*O*_):

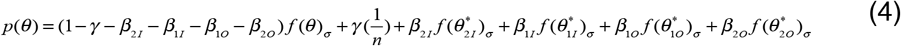

where *β*_2*I*_ is the probability of misreporting the second inner flanker (innermost) as the target and *β*_2*O*_ is the probability of misreporting the second outer (outermost) flanker as the target.

We fit the models to the individual data using the MemToolBox^54^. We compared models’ performance by comparing means Akaike information criterion with correction (AICc) to the individual data.

For each fitted model, we calculated target reporting rate (*P_T_*) by subtracting the accumulative guessing rate and misreport rate from 1, such that *P_T_* = 1− *γ*, *P_T_* = 1− *γ* − *β*,. *P_T_* = 1− *γ* − *β*_1*I*_ − *β*_1*O*_, *P_T_* = 1− *γ* − *β*_1*I*_ − *β*_1*O*_ − *β*_2*I*_ − *β*_2*O*_, for the standard mixture, misreport, two-misreport and four-misreport, respectively.

### Results

**Figure 2A** plots the error distribution for each condition. We first examined the bias of the errors by calculating mean error for each subject in each condition. Mean error was close to zero in the uncrowded condition (M = −1.17, SD = 1.62), the two-flanker condition (M = −1.33, SD = 2.46) and the four-flanker condition (M = −1.16, SD = 2.7). For each observer in each condition, we calculated precision as the inverse of the variance of the errors (**Fig. 2B**). We conducted a one-way Analysis of Variance (ANOVA) on precision with crowding conditions (uncrowded vs. two-flanker vs. four-flanker) as a within subject factor. A main effect on precision was observed, *F*(2,24) = 28.05, *p* < 0.001, partial *η^2^* = 0.70, indicating higher precision in the uncrowded condition than in the two-flanker condition, *t*(12) = 5.30, *p* < 0.001, Cohen’s *d* = 1.93, and in the two-flanker condition compared to the four-flanker condition, *t*(12) = 2.18, *p* = 0.050, Cohen’s *d* = 0.56.

**Figure 2.**
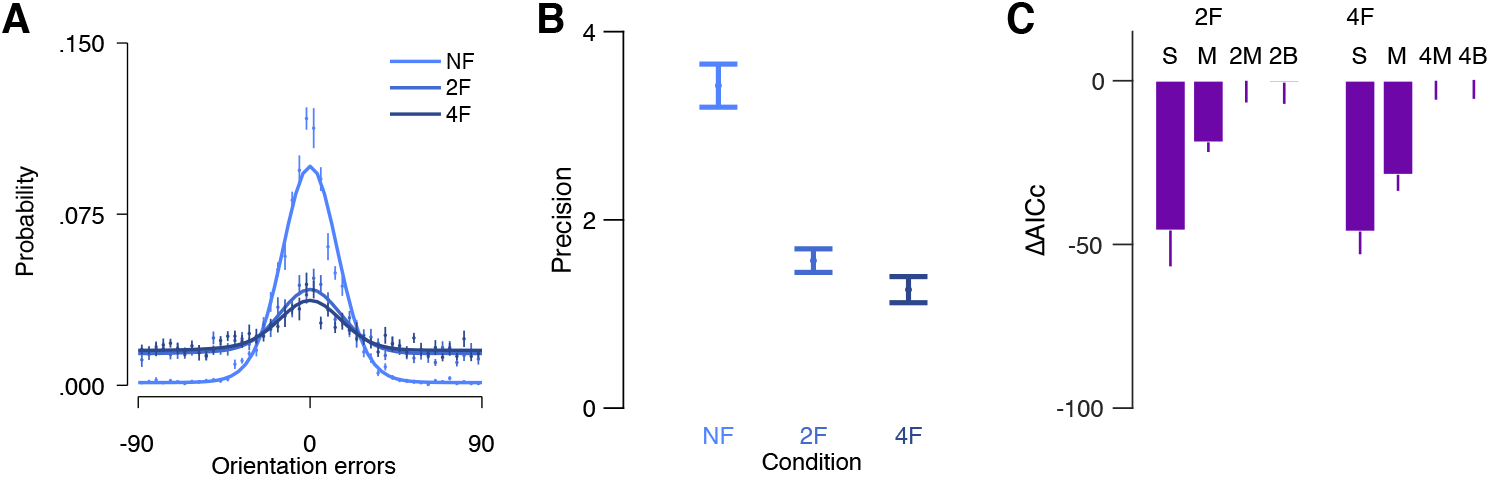
Error distribution and model comparison of Experiment 1. (**A**) Mean errors for each of the three crowding conditions. Solid lines are the best performing model in each crowding condition. (**B**) Mean precision (inversed SD in radiance) of the errors in each crowding condition. (**C**) Mean AICcs for each model subtracted from the AICc of the best performing model (AICc_2M_ – AICc_i_ and AICc_4M_ – AICc_i_, in the two-flanker and four-flanker respectively) in each crowded condition. UC: uncrowded, 2F: two-flanker and 4F: four-flanker. Models: S: standard mixture, M: misreport, 2M: two-misreport, 2B two-misreport with bias, 4M: four-misreport, 4B four-misreport with bias. Error bars are within subject ±1 SEM.

### Probabilistic models

In uncrowded conditions, the standard mixture model described well the distribution of the errors (**Fig. 2A**). For each crowding condition we chose the best model among two relevant models by calculating the Akaike information criterion with correction (AICc) for each observer. **Figure 2C** shows mean AICc for the relevant models in each flanker condition. In both crowded conditions, the model with multiple misreports outperformed the other two models. First, the standard misreport model outperformed the standard mixture model in the two-flanker and the four-flanker conditions, *t*(12) = 4.63, *p* = 0.001, Cohen’s *d* = 0.72, and *t*(12) = 2.57, *p* = 0.025, Cohen’s *d* = 0.64 respectively. In the two-flanker condition, the two-misreport model outperformed the standard misreport model, *t*(12) = 4.54, *p* = 0.001, Cohen’s *d* = 0.77. In the four-flanker condition, the four-misreport outperformed the standard misreport model, *t*(12) = 2.58, *p* = 0.024, Cohen’s *d* = 0.56. These results suggest that in crowded conditions observers misreported specific flankers more than others.

To check for averaging errors in the inner-outer asymmetry, we tested whether observers averaged the target with the outer flanker by calculating a mean bias toward the outer flanker (**Supplementary Information**). To do this, we realigned the error directions so that the first-outer flanker was always positive with respect to the target. Calculation of the mean of the errors revealed a positive mean bias—i.e. mean errors were shifted towards the outer flanker (**Figure SI1**). However, model comparisons revealed no advantage for models with bias compared to models without bias (**Figure 2C**), suggesting that heavier weights over the outer flanker (sampling bias) in the two- and four-misreport models, without an additional averaging process, is sufficient to explain the mean bias.

Next, we analyzed the fitted parameters of the best performing model in each condition. **Figure 3** depicts the mean fitted variability (σ) guessing rate (γ) and target reporting rate (*P_T_*) of the best fitted models in each condition. To assess the effect of crowding on performance, we conducted one-way, repeated measure analyses of variance (ANOVA) on variability, guessing rate and target reporting rate as dependent variables, with crowding condition as a within subject factor. There was a main effect of crowding condition on guessing rate, F(2,24) = 5.78, *p* = 0.009, partial *η^2^* = 0.33, but not on variability, F(2,24) = 2.37, *p* = 0.115. There was a main effect of crowding condition on target reporting rate, F(2,24) = 71.21, *p* < 0.001, partial *η^2^* = 0.86, revealing higher probability of reporting on the target under the uncrowded condition compared to the two-flanker condition, *t*(12) = 8.27, *p* < 0.001, Cohen’s *d* = 2.93, and the four-flanker condition, *t*(12) = 12.35, *p* < 0.001, Cohen’s *d* = 4.74. The probability of reporting on the target under the two crowded conditions did not differ significantly, *t*(12) = −1.40, *p* = 0.188.

**Figure 3.**
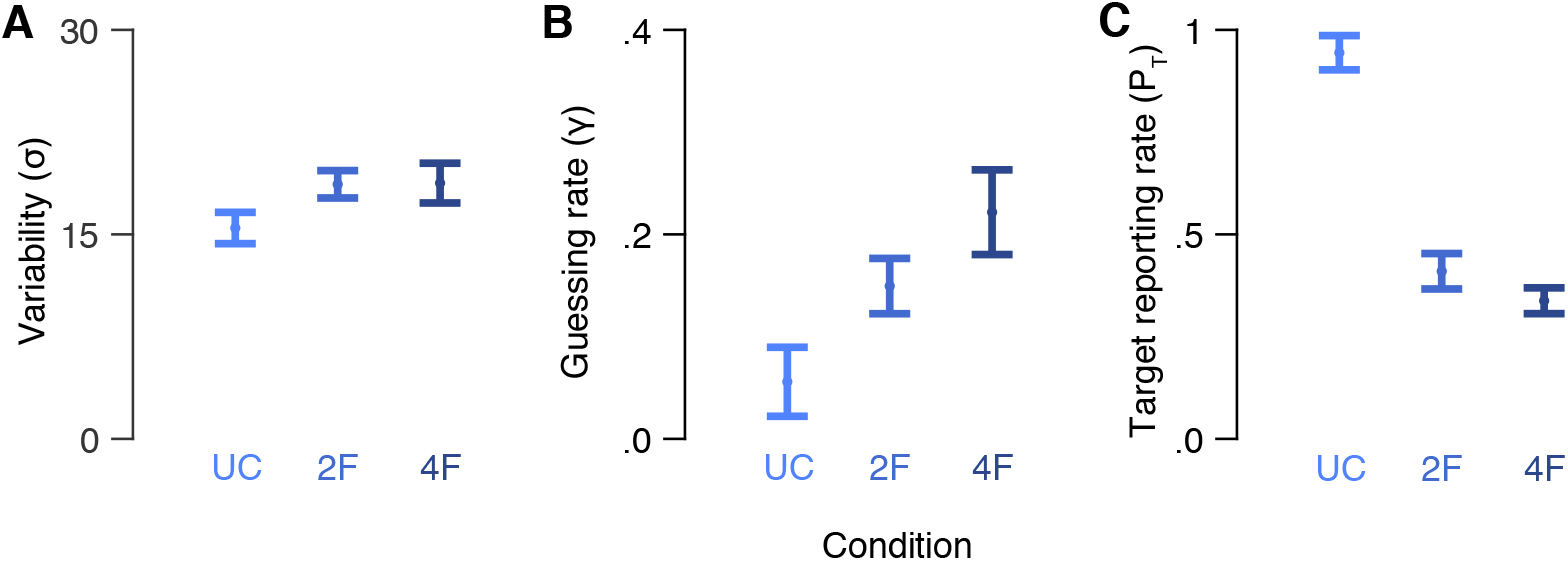
Model parameters in Experiment 1. (**A**) Mean variability (σ), (**B**) guessing rate (γ) and (**C**) target reporting rate (P_T_) for each crowding condition. UC: uncrowded, 2F: two-flanker, 4F: four-flanker. Error bars are ±1 SEM.

In order to test the contribution of each flanker to crowding errors, we compared the misreport rates of the different flankers. **Figure 4** depicts the distributions of reports around the value of each flanker and the probability of reporting each presented item. Importantly, in the two-misreport model (the two-flanker condition), the misreport rate of the outer flanker was significantly higher than the misreport rate of the inner one, *t*(12) = −2.59, *p* = 0.024, Cohen’s *d* = 1.24. In the four-misreport model (four-flanker condition), we conducted a one-way ANOVA on misreport rate with flanker positions (second inner, first inner, first outer and second outer) as within subject factor. Flanker position had a significant effect on misreport rate, *F*(3,36) = 5.74, *p* = 0.003, partial *η^2^* = 0.32, indicating that misreport rate differed across flankers. Planned comparisons revealed that the misreport rate of the first (adjacent) outer flanker (M= 0.24, SD = 0.13), was higher compared to the averaged probability of reporting on the other three flankers (M= 0.07, SD = 0.07), *t*(12) = 3.43, *p* = 0.005, Cohen’s *d* = 1.59. These findings support the view that the inner-outer asymmetry is due to misreport of the first (adjacent) outer flanker as the target.

**Figure 4.**
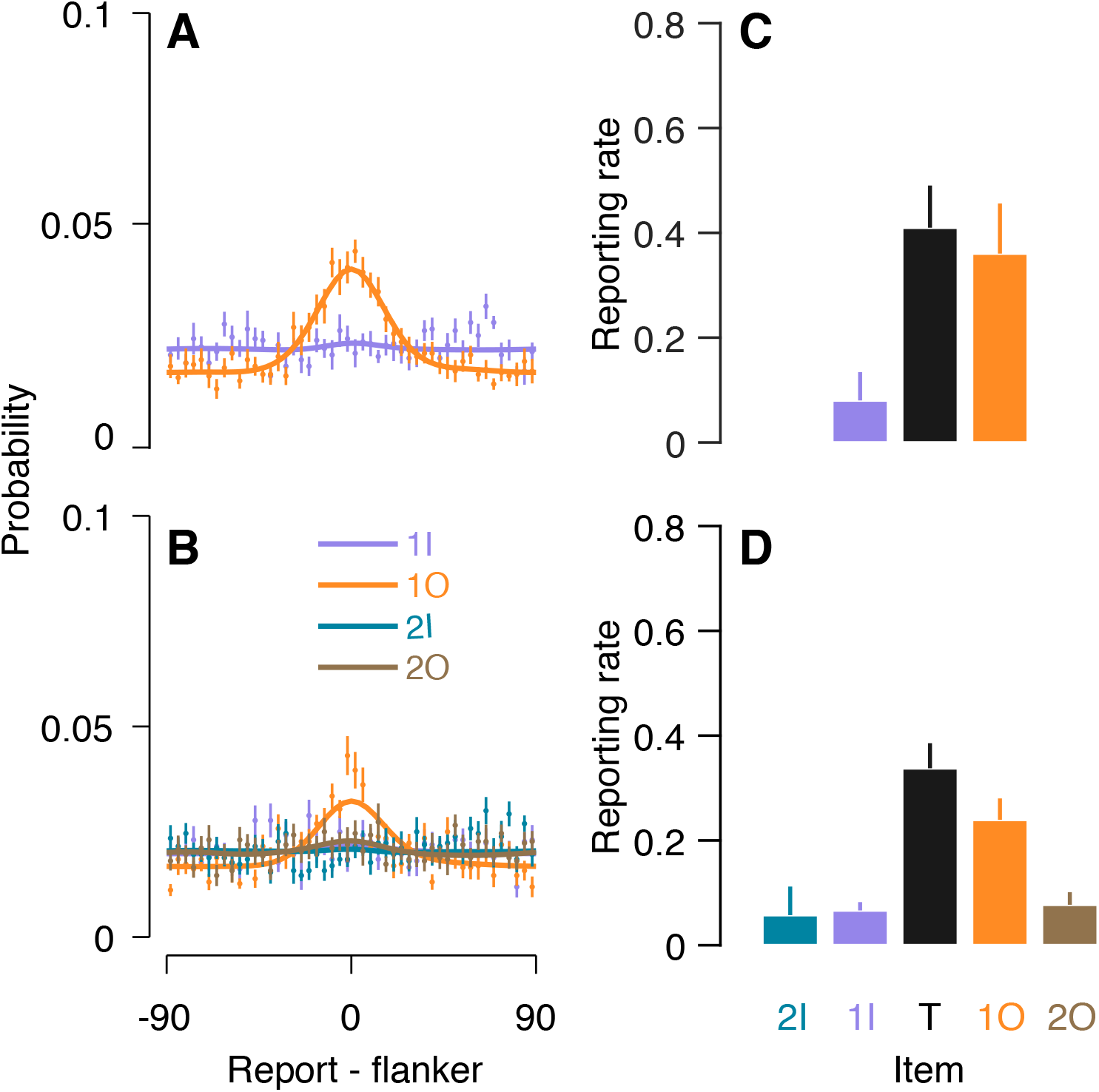
Misreporting the outer flanker as the target in Experiment 1. (**A** and **B**) Distribution of reports around the value of each flanker (0) for the (**A**) two-flanker and (**B**) four-flanker conditions, along with the fitting (solid lines) of the two-misreport model and the four-misreport model respectively. (**C** and **D**) Reporting rates for each item in the two-flanker (**C**) and four-flanker (**D**) conditions. Flanker report rates are the fitted independent β to each flanker in the two-misreport (**C**) and the four-misreport (σ) (**D**) models. 2I: 2nd inner flanker, 1I: 1st inner flanker, T: target, 1O: 1^st^ outer flanker, 2O: 2^nd^ outer flanker. Error bars are within subject ±1 SEM.

The results of Experiment 1 show that a reduction in overall precision of orientation estimation tasks under crowding is due to reporting a flanker instead of the target rather than an increase in variability (σ) over the representation of the target. This finding replicate that of Yashar et al (2019). Importantly, the results show that when the target is flanked radially from both sides, observers often misreport the orientation of the outer flanker that is adjacent to the target instead of the target. Observers misreported a flanker that was inner or not adjacent to the target in a much smaller proportion of the trials.

## Experiment 2

Following the results of Experiment 1, in Experiment 2 we examined whether observers’ tendency to report the outward item (a flanker) instead of the central one (the target) is a mutual process. If this is indeed a mutual process, when the target is the outward item observers will often misreport the central item (a flanker) as the target. We used the same display as in Experiment 1, except that now the target was shifted one Gabor outward. That is, in the two-flanker condition, the target was the outermost Gabor, and, in the four-flanker condition, the target was flanked by the outermost Gabor (see Fig. 1C). Note that the target eccentricity was increased in Experiment 2 compared to Experiment 1. Thus, if crowding zone size merely scales with eccentricity, we predict stronger crowding in Experiment 2 compared to 1 in all crowded displays. Alternatively, if the crowding zone is also skewed outward – that is, with more interference from the outer flanker—then we predict a substantial reduction in crowding in the two-flanker condition in which the outmost item is the target, and a strong crowding effect in the four-flanker condition, in which the target is flanked by an outer flanker.

### Method

#### Observers

Fourteen undergraduate and graduate students (10 females: age range 19 to 36 years old, M=25.71, SD=5.03), from The University of Haifa participated in this experiment for a course credit or a 40 ILS per hour monetary payment (around $12). All observers were naive to the purposes of the experiment. All observers reported having normal or corrected-to-normal visual acuity and normal colour vision, and none reported attention deficits or epilepsy. The observers signed a written informed consent form before the experiment. The experimental procedures were approved by The University Committee on Activities Involving Human Subjects at The University of Haifa (No. 373/18).

#### Apparatus

The apparatus was the same as described in Experiment 1.

#### Stimuli and procedure

**Figure 1A** and **C** illustrate a trial sequence and stimulus condition in Experiment 2. Stimuli and procedure were the same as in Experiment 1, except for the location of the target. In this experiment, the target appeared on the horizontal meridian with 8.5° eccentricity, either in the right or left hemifield. In the two-flanker condition, the target was the outer item, whereas in the four-flanker condition, the target was located near the outer flanker. Here too, the centre-to-centre distance between two adjacent items was 1.5°. In the two-flanker condition, flanker eccentricities were 5.5° and 7° for the second inner (2I) and first inner (1I) flankers respectively. In the four-flanker condition, flanker eccentricities were 4°, 5.5°, 7° and 10° for the third inner (3I), second inner (2I), first inner (1I) and first outer (1O) flankers respectively.

#### Design

The design was the same as described in Experiment 1.

##### Models and analyses

The statistical analyses were the same as described in Experiment 1.

### Results

**Figure 5A** illustrates the error distribution for each condition. Here again, we first analyzed the bias of the errors by calculating mean error for each subject in each condition. Mean error was close to zero in the uncrowded condition (M= −1.01, SD = 2.29), the two-flanker condition (M = −0.41, SD = 2.75) and the four-flanker condition (M= 0.34, SD = 4.03). As in Experiment 1, we calculated precision as the inverse of the variance of the errors for each observer in each condition (**Fig. 5B**). One-way ANOVA analyses on precision and crowding condition as within subjects factor revealed a significant main effect of crowding conditions on precision, F(2,26) = 44.30, *p* = 0.000, partial *η^2^* = 0.77, indicating higher precision in the uncrowded condition compared to the two-flanker condition, *t*(13) = 6.13, *p* = 0.000, Cohen’s *d* = 1.45, and in the two-flanker condition compared to the four-flanker condition, *t*(13) = 4.19, *p* = 0.001, Cohen’s *d* = 1.60.

**Figure 5.**
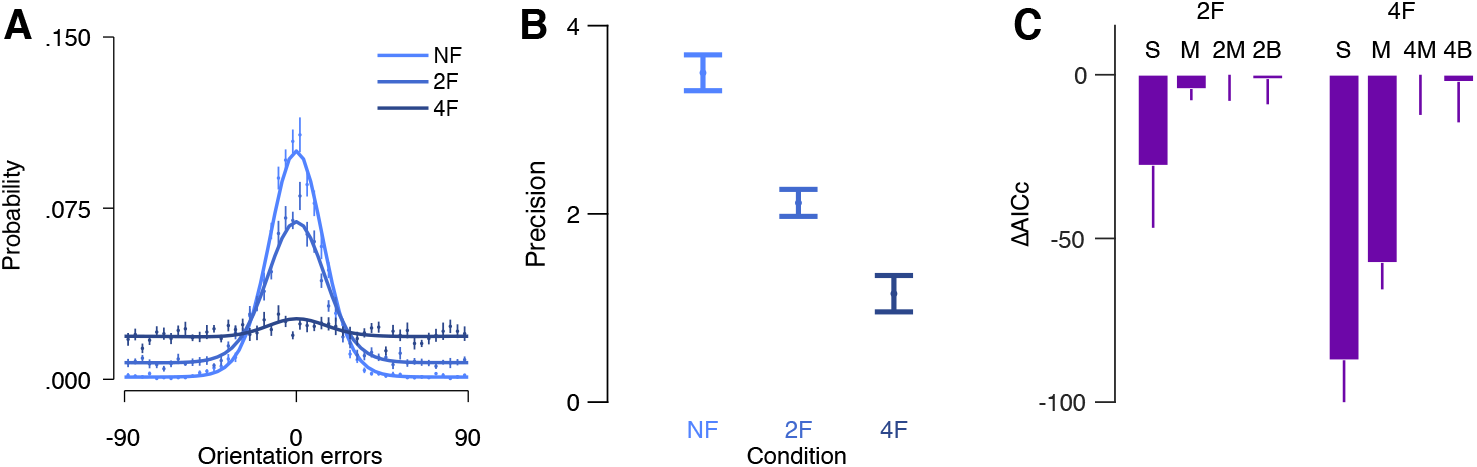
Error distributions and model comparisons of Experiment 2. (**A**) Mean errors for each of the three crowding conditions. Solid lines represent the best performing model in each crowding condition. (**B**) Mean precision (inverse of the SD of the errors in radiance) of the errors in each crowding condition. (**C**) Mean AICcs for each model subtracted from the AICc of the best performing model in each crowded condition (the two-misreport model and the four-misreport model in the two-flanker and the four-flanker conditions, respectively). UC: uncrowded, 2F: two-flanker and 4F: four-flanker. Models: S: standard mixture, M: misreport, 2M: two-misreport, 2B two-misreport with bias, 4M: four-misreport, 4B four-misreport with bias. Error bars are within subject ±1 SEM.

### Probabilistic models

In the uncrowded condition, the standard mixture model described well the distribution of the errors **(Fig. 5A**). For each flanker condition, we calculated the Akaike information criterion with correction (AICc) for each observer, in order to choose the best model among the two relevant models. **Figure 5C** presents mean AICc for the relevant models in each crowding condition. In the two-flanker condition, there was no best-fitting model, meaning that no significant differences were found between the standard misreport model and the standard mixture model, *t*(13) = 1.30, *p* = 0.216 and between the independent two-misreport model and the standard misreport model, *t*(13) = 1.25, *p* = 0.232. This result is explained by small crowding interference in this condition. However, in the four-flanker condition, the standard misreport model outperformed the standard mixture model, *t*(13) = 4.06, *p* = 0.001, Cohen’s *d* = 0.69, and the independent four-misreport outperformed the standard misreport model, *t*(13) = 3.80, *p* = 0.002, Cohen’s *d* = 0.91. In this condition, misreport rates were higher for some flankers compared to others. As in Experiment 1, we tested the fitting of models with mean bias towards the outer flanker (**Supplementary Analysis**). Again, adding the mean bias component did not improve model fitting (**Figure 5C and Figure SI2**), which indicates that the misreport models are sufficient to explain our results. Next, we analyzed the fitted parameters of the best performing model (smallest AICc) in each condition, model two-misreport and four-misreport respectively. **Figure 6** depicts the mean fitted variabilities (σ) and guess rates (γ) of the best fitted models in each condition. In order to examine the effect of crowding on performance, we conducted a one-way ANOVA with crowding condition (uncrowded vs. two-flanker vs. four-flanker) as a within subject factor and variability (σ), guess rate (γ) and target reporting rate (*P_T_*) as dependent variables. Significant differences were found between the crowding conditions for γ, F(2,26) = 9.86, *p* = 0.001, partial *η^2^* = 0.43,, F(2,26) = 15.71, *p* = 0.001, partial *η^2^* = 0.55, and *P_T_*, F(2,26) = 71.66, *p* < 0.001, partial *η^2^* = 0.85. Specifically, the probability to report on the target decreased as the number of flankers increased, with significant differences between the uncrowded condition and the crowded conditions (two-flanker: *t*(13) = 3.76, *p* = 0.002, Cohen’s *d* = 1.35, four-flanker: *t*(13) = 21.17, *p* < 0.001, Cohen’s *d* = 7.17), and between the two crowded conditions, *t*(13) = 6.36, *p* = 0.001, Cohen’s *d* = 2.26.

**Figure 6.**
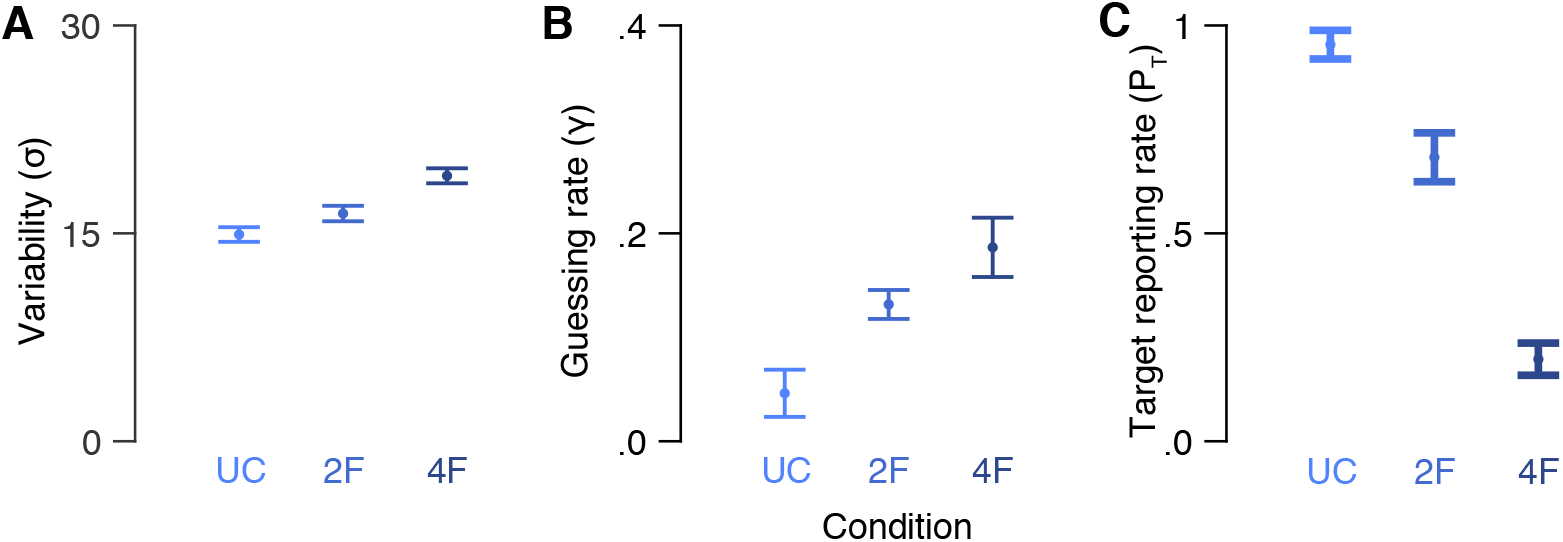
Model parameters in Experiment 2. (**A**) Mean standard deviation (σ), (**B**) guessing rate (γ) and (**C**) target reporting rate (P_T_) for each crowding condition. Model parameters are of the S, M2 and M4 models in the uncrowded, two-flanker and four-flanker conditions, respectively. UC: uncrowded, 2F: two-flanker and 4F: four-flanker. Error bars are ±1 SEM.

We assessed the contribution of each flanker to crowding errors by comparing the misreport rates of the different flankers. **Figure 7** depicts the report distribution around the value of each flanker and the probability of reporting each presented item. In the two-misreport model (the two-flanker condition), in which subjects were asked to report on the outer flanker, we found insignificant effect for flankers’ positions, (t(13) = −0.35, *p* = 0.730). In the four-misreport model (four-flanker condition), we conducted a one-way ANOVA on misreport rate with flanker position (third inner, second inner, first inner, and first outer) as within subject factor. A significant effect for flanker position was obtained in the four-flanker condition in which the target was located near the outer flanker, F(3,39) = 12.94, *p* = 0.000, partial *η^2^* = 0.50. Here, planned comparisons revealed that the misreport rate of the first (adjacent) outer flanker (1O) (M = 0.44, SD = 0.25) was significantly higher compared to the averaged misreport rate of the other flankers (M= 0.06, SD = 0.09), *t*(13) = 4.41, *p* = 0.001, Cohen’s *d* = 2.03. These findings are consistent with Experiment’s 1 results, and support the view that observers often misreport the first (adjacent) outer flanker as the target.

**Figure 7.**
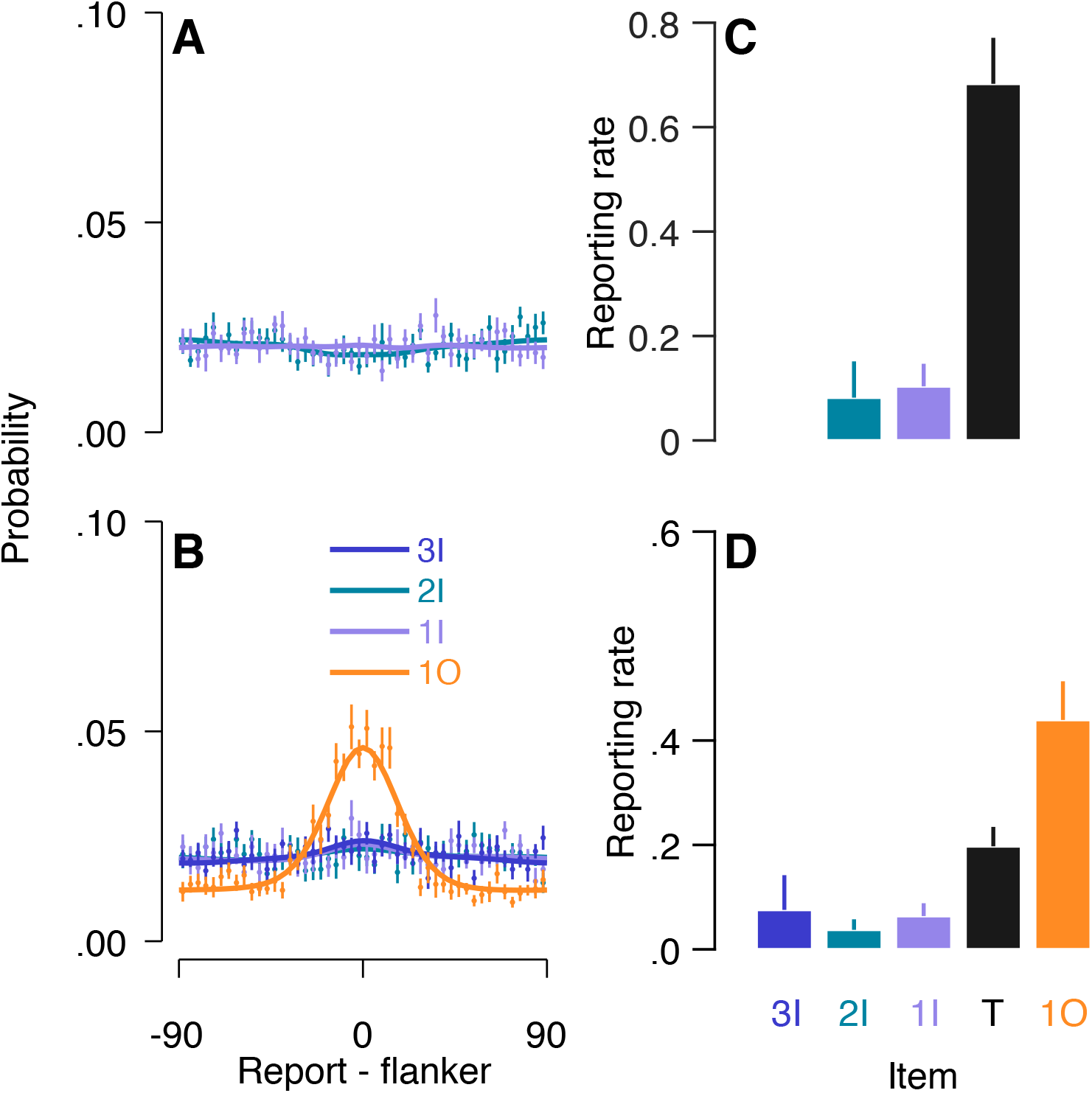
Misreporting the outer flanker as the target in Experiment 2. (**A** and **B**) Distribution of reports around the value of each flanker (0) for the (**A**) two-flanker and (B) four-flanker conditions, along with the fitting (solid lines) of the two-misreport and the four-misreport models respectively. (**C** and **D**) Reporting rates for each item in the two-flanker (**C**) and four-flanker (**D**) conditions. Flanker report rates are the fitted independent β to each flanker in the two-misreport (**C**) and the four-misreport (**D**) models. 3I: 3^rd^ Inner flanker, 2I: 2^nd^ Inner flanker, 1I: 1^st^ Inner flanker, T: target, 1O: 1^st^ outer flanker. Error bars are within subject ±1 SEM.

The results of Experiment 2 show that the reduction of overall precision in orientation estimation task under crowding is due to both reporting a flanker instead of the target and increased variability (σ) over the representation of the target. Here again, we found that observers reported the adjacent flanker that was more eccentric rather than the target. However, when the target was the outer item, crowding was reduced and there was no significant difference in the misreport rate (*P_T_*) between the flankers. Furthermore, as shown in previous studies^6,55^, in both Experiment 1 and Experiment 2 we found that the frequency of responses around the target decreases with an increase in the number of flankers.

Finally, we examined the effect of flanker position on crowding interference. To do so, we conducted an independent t-test for each crowded condition and compared the target reporting rate (*P_T_*) and standard deviation (σ) between Experiment 1 and Experiment 2. *P_T_* analyses revealed that in the uncrowded condition, target reporting rate did not differ significantly, *t*(25) = −0.39, *p* = 0.701, meaning that observers were unaffected by item eccentricity per se. However, significant differences were found between the two experiments in the two crowded conditions. In the two-flanker condition, when observers reported the outer item (Experiment 2) crowding was smaller than when observers reported the item in the middle (Experiment 1), *t*(25) = −2.69, *p* = 0.013, Cohen’s *d* = 1.04. Although we increased the target eccentricity, crowding interference was reduced when observers reported the outer item (Experiment 2), suggesting that our findings cannot be explained by target eccentricity alone. In the four-flanker condition, when the outer target was flanked by an outer flanker (Experiment 2), crowding was stronger than when the central target was flanked by two outer flankers (Experiment 1), *t*(25) = 2.40, *p* = 0.024, Cohen’s *d* = 0.92 (see^56^ for a similar finding). Although the four-flanker condition in Experiment 2 might involve a higher degree of uncertainty compared to the four-flanker condition in Experiment 1, the fact that observers misreported the adjacent outer flanker here too is consist with the observed findings in Experiment 1. Finally, no differences were observed for σ between Experiment 1 and Experiment 2, with t-values bigger than 0.30 in each crowding condition.

## Discussion

Using an orientation estimation task and probabilistic models designed to assess the contribution of each flanker to crowding errors, we showed that the hallmark inner-outer asymmetry in crowding reflects misreports of the outer flanker as the target. First, as in previous studies^18,44,51^, our results showed that orientation crowding errors reflect reporting a flanker instead of the target (misreport errors). Second, we found that in a radial crowding display (Experiment 1), in which a target is flanked on both sides, the misreport rate of the outer flanker was significantly higher compared with the misreport rate of the inner one. Third, when two flankers appeared on each side of the target (four-flanker condition), the misreport rate of the outer flanker adjacent to the target was higher than the misreport rate of the most eccentric flanker. Finally, when the target was the outer item (Experiment 2, two-flanker condition) crowding errors were substantially reduced.

Our misreport mixture models provides a stochastic approximation of previously proposed population coding models^39,42,51^. These models explain crowding by a weighted summation of population coding within a receptive field. These models rely on an integration process, and hence they can be considered pooling models^50^. Here, the misreport model with independent flanker components outperformed models with either no misreport component or a single misreport component to all flankers. Thus, our modeling explains misreport errors due to pooling over feature representations.

Simple pooling models of crowding predict averaging of target and flanker features ^38,41^. Within the context of the inner-outer asymmetry, simple pooling models assert weighted average with more weight to the outer flanker and, therefore, predict mean bias towards the outer flanker^33^. Here, we show that adding a mean bias as a free parameter to the misreport model does not improve model fitting. This finding is inconsistent with previous accounts that suggests that crowding errors reflect a combination of misreport and averaging errors^43,52^. Our study, therefore, supports a single pooling mechanism account of crowding.

Importantly, our results show that the inner-outer asymmetry can be explained by a weighted summation in which the peripheral (outer) item receives heavier weight than the foveal (inner) item. In terms of population coding models, heavier weight over the peripheral item perhaps can be implemented by assuming that summation weights scale with receptive field size, such that heavier weights are assigned to more peripheral items (larger receptive field). A similar principle was proposed with a Bayesian inference framework and with some assumption regarding the growth of receptive field size in the visual periphery^27^. In particular, the number of receptive fields that cover only the outer item is larger than the number of receptive fields that cover only the inner item. Here we suggest that larger receptive fields in the periphery render errors towards the outer flanker.

Our results are inconsistent with the cortical magnification account. According to this view, crowding asymmetry results from cortical mapping in which the outer flanker is closer to the target than the inner one^6,37^. This explanation predicts that within the same display (i.e. same cortical distances), crowding interference will be the same regardless of whether observers are reporting on the middle or the outer item. Thus, the reduction of crowding interference in the two-flanker condition of Experiment 2 compared to Experiment 1 suggests that cortical magnification alone is insufficient to explain the inner-outer asymmetry of crowding^27^.

In this study we investigated the inner-outer asymmetry along the horizontal meridian by using a symmetrical display. Although previous studies found the asymmetry effect along the horizontal, but not along the vertical meridian^24^, it is still unclear whether and how this effect is reflected in a typical symmetrical crowding display across the visual field. Furthermore, our results are limited to orientation errors involved in crowding asymmetry. Since recent studies found dissociation of crowding errors across dimensions, such as orientation, color and spatial frequency^51^, and between color and motion^57^, further work is needed to determine whether the current findings apply to other feature dimensions. Moreover, the use of Gabor patches also limited our ability to distinguish between object level and feature level. Due to previous studies that pointed to different processes involved at these levels^34,51,58^, further work should use a different type of stimulus, which would allow a clear distinction between those levels. Finally, although our findings support population based pooling models, they do not reject simple pooling models of crowding. To do so, future studies should directly test the fitting of an averaging model (a weighted average between target and flankers).

## Conclusions

Our findings reveal that in a typical radial crowding display, observers confuse the target with the nearby outer flanker, but not vice versa. This confusion occurs regardless of the number of flankers and reported item position. Probabilistic mixture models explain that these results as due to heavier weights toward the visual periphery. These findings are consistent with recent models that explain crowding in terms of population coding with weighted summation within receptive fields. Our results demonstrate a counterintuitive phenomenon: within a region of selection, the more eccentric item dominates appearance.

## Data availability

The data and analysis codes are available from the corresponding author upon request.

## Acknowledgments

We would like to thank Hadas Nathan Gamliel for her invaluable help with data collection.

## Author Contributions

AY conceptualized this study. AS collected the data. AS and AY contributed to the research design, statistical analysis, interpretation of the data and writing.

## Additional information

Correspondence and requests for materials should be addressed to Amit Yashar.

