## Supplementary Information for "Mixture model investigation of the inner-outer asymmetry in visual crowding reveals a heavier weight towards the visual periphery"

### Supplementary Analysis

To test the bias of the errors towards either the inner or outer flanker, we aligned the direction of errors such that the value of the first outer flanker was always positive in respect to the target (0). Hence, a positive mean bias will indicate averaging errors with the first outer flanker. In Experiment 2, two-flanker condition, we aligned the direction of the errors toward the first inner flanker since in that condition there was no outer flanker.

*The two-misreport with bias mode*l and the *four-misreport with bias mode*l (Equation SI.1 and SI.2 respectively) have a free parameter for the mean of the distribution around the target. These models have five and seven free parameters, respectively:

$$p(\theta) = (1-\gamma-\beta_{1I}-\beta_{1O})f(\theta)_{\mu,\sigma} + \gamma(\frac{1}{n}) + \beta_{1I}f(\theta^*_{1I})_\sigma + \beta_{1O}f(\theta^*_{1O})_\sigma \qquad \text{(SI.1)}$$

$$p(\theta) = (1-\gamma-\beta_{2I}-\beta_{1I}-\beta_{1O}-\beta_{2O})f(\theta)_{\mu,\sigma} + \gamma(\frac{1}{n}) + \beta_{2I}f(\theta^*_{2I})_\sigma + \beta_{1I}f(\theta^*_{1I})_\sigma + \beta_{1O}f(\theta^*_{1O})_\sigma + \beta_{2O}f(\theta^*_{2O})_\sigma \qquad \text{(SI.2)}$$

In both models, $\mu$ is mean bias of the distribution around the estimated value.

### Supplementary Results

Fitting the models with bias did not change the overall pattern of results; misreport rates, rather than mean bias shifts, explain the inner outer asymmetry In both Experiment 1 and Experiment 2 (Figure SI1 and SI2 respectively),

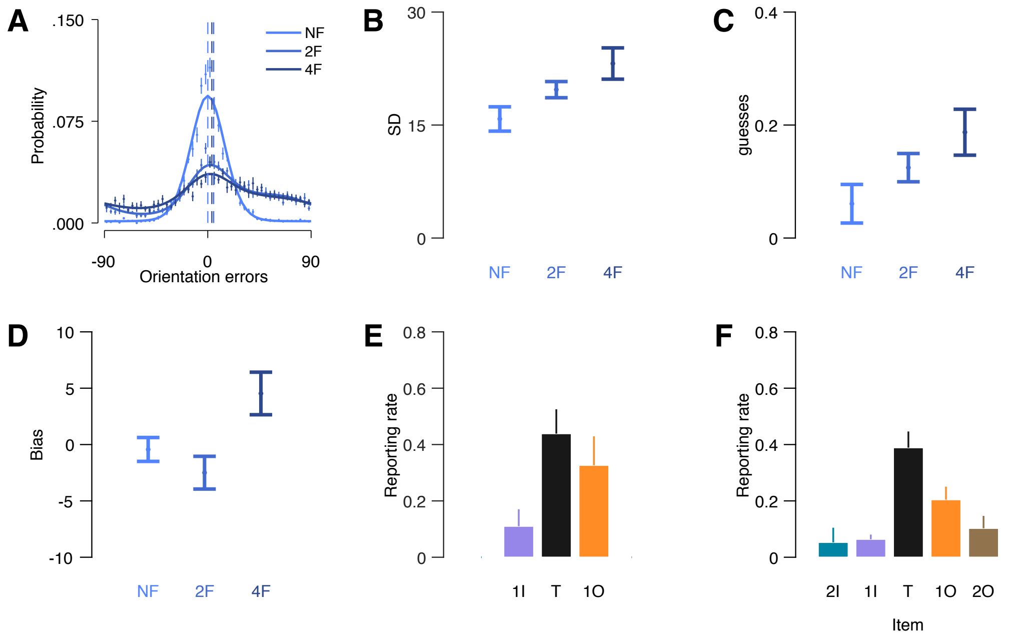


**Figure SI1**. **Experiment 1 data and fitted parameters of the models with bias.** Error distribution and model comparison of Experiment 1 data realigned so that the first outer flanker is positive in respect to the target. (**A**) Mean errors for each of the three crowding conditions. Solid lines are the best performing model in each crowding condition. Dashed line are error means. (B-F) The free parameters of the tow-misreport with bias and the four misreport with bias in each crowding condition:  (**B**) Mean SD, (**C**) Mean guesses, (**D**) Mean bias, (**E**) Mean misreport rate in the two-flanker condition, (**F**) Mean misreport rate in the four-flanker condition. Error bars are within subject ±1 SEM. UC: uncrowded, 2F: two-flanker and 4F: four-flanker. 2I: 2nd inner flanker, 1I: 1st inner flanker, T: target, 1O: 1st outer flanker, 2O: 2nd outer flanker.

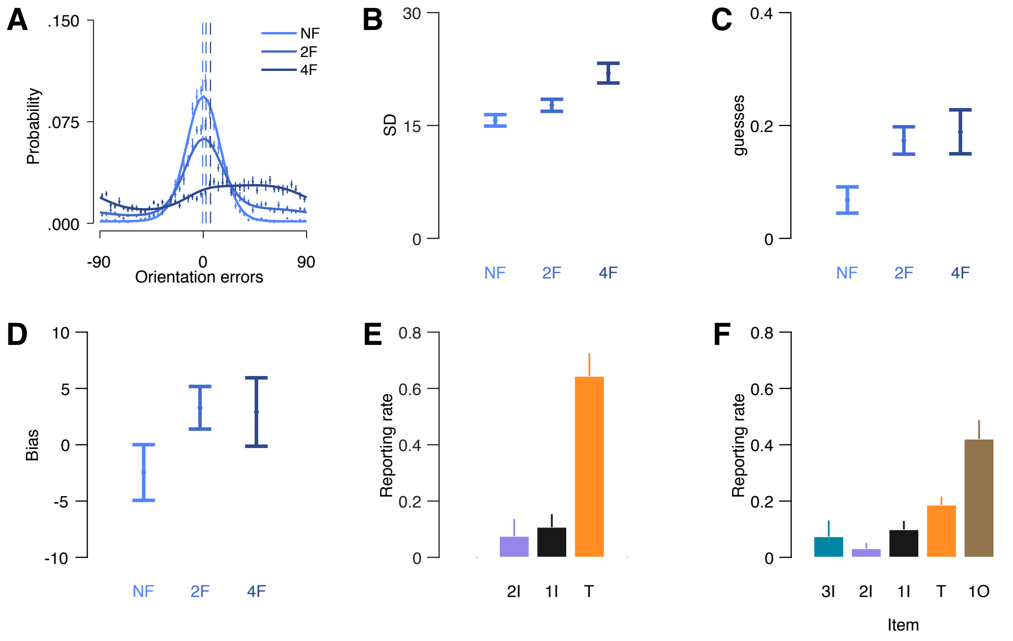


**Figure SI2**. **Experiment 2 data and fitted parameters of the models with bias**. Error distribution and model comparison of Experiment 1 data realigned so that the first outer flanker is positive in respect to the target. (**A**) Mean errors for each of the three crowding conditions. Solid lines are the best performing model in each crowding condition. Dashed line are error means. (B-F) The free parameters of the tow-misreport with bias and the four misreport with bias in each crowding condition: (**B**) Mean SD, (**C**) Mean guesses, (**D**) Mean bias, (**E**) Mean misreport rate In the two-flanker condition, (**F**) Mean misreport rate in the four-flanker condition. Error bars are within subject ±1 SEM. UC: uncrowded, 2F: two-flanker and 4F: four-flanker. 2I: 2nd inner flanker, 1I: 1st inner flanker, T: target, 1O: 1st outer flanker, 2O: 2nd outer flanker.
